# Atypical Alpha Oscillatory EEG Dynamics in Children with Angelman Syndrome

**DOI:** 10.1101/2025.04.03.647110

**Authors:** Abigail H. Dickinson, M. Sapphire Bowen-Kauth, Jeremy J. Shide, Anna E. Youngkin, Nishitha S. Hosamane, Courtney A. McNair, Declan P. Ryan, Catherine J. Chu, Michael S. Sidorov

## Abstract

**Objectives:** Biomarkers of atypical brain development are crucial for advancing clinical trials and guiding therapeutic interventions in Angelman syndrome (AS). Electroencephalography (EEG) captures well-characterized developmental changes in peak alpha frequency (PAF) that reflect underlying neural circuit maturation and may provide a sensitive metric for mapping atypical neural trajectories in AS.

**Method:** We analyzed EEG recordings from 159 children with AS (ages 1–15 years) and 185 age-matched typically developing (TD) controls. PAF was quantified using a well-established curve-fitting method applied to 1/f-corrected power spectra. To validate robustness, we further evaluated PAF using an alternative prominence-based peak detection approach across varying detection thresholds.

**Results:** Significant disruptions in PAF were evident in children with AS. While over 90% of EEGs from TD children exhibited a clear alpha peak, fewer than 50% of EEGs from children with AS showed a detectable PAF. Furthermore, when PAF was present, its frequency was significantly lower in AS children and did not show the typical age-related increases observed in TD children. Validation analyses confirmed consistently lower rates of PAF detection in AS across varying sensitivity thresholds, demonstrating the robustness of these results.

**Conclusions:** PAF is a robust and developmentally sensitive marker of disrupted neural maturation in children with Angelman syndrome. As a quantifiable and sensitive measure of neural disruptions in AS, PAF has the potential to complement and enhance existing clinical trial outcome assessments by providing an objective index of underlying brain function. Future analyses will explore individual differences related to PAF in AS, to better understand mechanistic insights to guide targeted therapeutic strategies.

## 1. INTRODUCTION

Angelman syndrome (AS) is a rare neurodevelopmental disorder characterized by significant cognitive and motor impairments, limited or absent speech, seizures, sleep disruptions, and pervasive developmental delays (Bird, 2014; Williams, 2005). These clinical features result from the loss of function of the maternal copy of the ubiquitin-protein ligase E3A (*UBE3A*) gene, typically due to deletions or mutations (Kishino et al., 1997). In neurons, the paternal *UBE3A* allele is silenced by the long, non-coding antisense transcript *UBE3A-ATS*, making the maternal allele the sole source of UBE3A protein (Meng et al., 2012; Rougeulle et al., 1998; Runte et al., 2001). The resulting UBE3A deficiency in AS disrupts neuronal and synaptic processes essential for early brain development, leading to the defining clinical characteristics of the disorder.

Preclinical research has leveraged the conserved paternal imprinting of UBE3A in mice (Albrecht et al., 1997; Judson et al., 2014) to explore treatments aimed at reactivating the paternal allele. These efforts have shown remarkable efficacy in AS mouse models (*Ube3a*^*m-/p+*^) (Huang et al., 2011; Meng et al., 2013; Meng et al., 2015; Milazzo et al., 2021; Vihma et al., 2024; Wolter et al., 2020), and have advanced to clinical trials (Elgersma and Sonzogni, 2021). Alongside preclinical progress in other mechanism-based therapies (Copping et al., 2021), these developments underscore the urgent need for robust biomarkers to evaluate and guide therapeutic interventions in AS (Berry-Kravis et al., 2018).

Electroencephalography (EEG) is a non-invasive, accessible, and scalable method for measuring brain activity that holds significant potential for identifying such markers. EEG provides a direct measure of brain activity and can capture subtle oscillatory dynamics targeted by disease-modifying therapies, offering a sensitive means to evaluate target engagement and therapeutic effects in real-time. EEG has been widely used to investigate markers of atypical neural function in neurodevelopmental disorders, including AS (Goodspeed et al., 2023). One of the most well-documented EEG findings in AS is an increase in low-frequency (∼1-4 Hz) delta rhythms (Bower and Jeavons, 1967; Boyd et al., 1988; Laan et al., 1997; Vendrame et al., 2012), an observation that has recently been supported by quantitative studies (Frohlich et al., 2019; Levin et al., 2022; Martinez et al., 2020; Sidorov et al., 2017). Increased delta in AS correlates with clinical severity (Hipp et al., 2021; Ostrowski et al., 2021), and is now being evaluated as a biomarker in clinical trials. However, while delta power is a crucial marker of generalized neural disruption, it may lack sensitivity to detect more nuanced changes in cortical circuit development or therapeutic gains.

Alpha oscillations (6–12 Hz) may serve as a complementary biomarker for mapping circuit-level changes in AS. These rhythms play a crucial role in neuronal timing, functional inhibition, and integrating information across brain regions (Foxe and Snyder, 2011; Jensen and Mazaheri, 2010; Klimesch et al., 2007) and exhibit well-established developmental changes that are closely linked to cortical circuit maturation and cognitive function (Freschl et al., 2022; Marshall et al., 2002). During typical development, a distinct peak in alpha frequencies (peak alpha frequency; PAF), emerges towards the end of the first year (∼6 Hz) and gradually increases in frequency throughout childhood, reflecting the maturation and growing efficiency of neural circuits (Freschl et al., 2002).

The developmental trajectory of alpha oscillations provides a valuable benchmark for identifying and tracking atypical brain development. For instance, deviations in PAF emergence and maturation have been observed in autism, Fragile X syndrome, and neurofibromatosis type 1 (Booth et al., 2023; Dickinson et al., 2018; Pedapati et al., 2022). Furthermore, developmental changes in PAF are correlated with non-verbal cognitive abilities (Carter Leno et al., 2021; Dickinson et al., 2019; Edgar et al., 2019), suggesting that PAF may serve as a direct index of the circuit-level changes underlying cognitive function—a core therapeutic target in AS clinical trials. Despite this potential, alpha oscillations in AS have not been rigorously analyzed across a broad developmental window. While prior studies have presented power spectra suggesting likely alpha alterations in AS (Frohlich et al., 2019; Ostrowski et al., 2021; Sidorov et al., 2017), it remains unclear whether alpha metrics, such as PAF, sensitively index disrupted network dynamics or exhibit an atypical age-related trajectory in this population.

This study addresses this gap by characterizing alpha dynamics, specifically power and PAF, in children with AS and comparing these metrics to an age-matched typically developing (TD) cohort. Additionally, we examine age-related changes in alpha dynamics across the two groups. By providing a detailed assessment of these metrics, we aim to evaluate their potential as biomarkers of brain development in AS. Given the extensive neural disruptions associated with AS, we hypothesize atypical alpha patterns, including reduced power and frequency, as well as deviations in age-related trajectories.

## 2. METHODS

### 2.1. Participants

This study utilized deidentified EEGs (n=159) from children with AS and typically developing (TD) controls (n=185), drawn from established repositories and previously published datasets (**Table 1**). Participants ranged in age from 0.9 to 14.9 years (TD: 5.8 ± 0.3 years; AS: 6.1 ± 0.3 years; **Fig. S1a**).

**Table 1:** Participant demographics. TD: typically developing, AS: Angelman syndrome.

| *Group* | *# of EEGs* | *Unique participants* | *Age (years)* | *Male* | *Female* | *Length of EEG (minutes)* |
| --- | --- | --- | --- | --- | --- | --- |
| TD | 185 | 185 | $5.8 \pm 0.3$ | 99 (54%) | 86 (46%) | $10.3 \pm 1.3$ |
| AS | 159 | 95 | $6.1 \pm 0.3$ | 102 (64%) | 57 (36%) | $24.7 \pm 2.1$ |

A total of 159 EEGs from individuals with AS were accessed through established repositories. Of these, 149 EEGs were collected as part of the AS Natural History Study (ClinicalTrials.gov identifier: NCT00296764) and shared via the LADDER database (Potter et al., 2024), and 10 were collected by the UNC Sleep Disorders Center (Levin et al., 2022). These recordings represented 95 unique participants, as 35 individuals contributed multiple recordings (**Supplementary Tables 1-2**). Genetic information was available for 89 of the 95 AS participants, with a genotype distribution representative of the general AS population (Bird, 2014): 21% class I deletion, 38% class II deletion, 3% atypical deletion, 6% deletion (unspecified), 10% paternal uniparental disomy, 4% imprinting defect, 15% UBE3A mutation, 3% abnormal DNA methylation (**Fig. S1b**).

Age-matched EEGs from TD controls were drawn from previously published datasets collected at three sites: Massachusetts General Hospital (MGH) (*n* = 98), UNC Sleep Disorders Center (*n* = 10), and UCLA (*n* = 77) (den Bakker et al., 2018; Dickinson et al., 2018; Dickinson et al., 2025; Levin et al., 2022; Ostrowski et al., 2021; Sidorov et al., 2017) (**Supplementary Table 3**). Children in these datasets were identified as TD by the studies through which their data were collected, based on the absence of neurodevelopmental or genetic conditions.

### 2.2. EEG processing

EEG data in the present analyses were derived from continuous (task-free) recordings pooled from multiple studies and sites. The specific system hardware configurations and recording parameters are detailed in **Supplementary Tables 2 & 3**. To address these variations, we applied data processing procedures designed to harmonize datasets and ensure consistent analysis across studies (Levin et al., 2022; Sidorov et al., 2017).

For clinical EEGs, periods of wake and sleep were identified and annotated by a trained neurologist, ensuring that only wakeful data were analyzed. Raw EEGs were preprocessed using methods adapted from prior studies (Levin et al., 2022; Sidorov et al., 2017) and implemented using custom MATLAB scripts.

Preprocessing included re-referencing to the common average, applying a 0.1 Hz high-pass filter, a 100 Hz low-pass filter, and a 60 Hz notch filter to remove noise and electrical interference.

Recordings were visually inspected to identify and exclude artifacts and excessively noisy channels. After artifact removal, an average of 17.0 ± 1.2 minutes of EEG data per recording underwent further analysis, with 10.3 ± 1.3 minutes for TD EEGs and 24.7 ± 2.1 minutes for AS EEGs (**Table 1**). All recordings retained at least 60 seconds of artifact-free data.

To ensure consistency, analyses focused on six specific channels representing three regions of interest: F3 and F4 (frontal), C3 and C4 (central), and O1 and O2 (occipital). Across all recordings (2064 total channels), 28 channels were excluded (1.35%), including 21 from AS recordings and seven from TD recordings. Missing channels affected 6.1% of participants (17 AS, 4 TD). If both channels for a region were missing, the region was excluded from analyses for that participant. This impacted 1.45% of participants (4 AS, 1 TD). Among these, four AS participants had one region excluded (3 frontal, 1 occipital), and one TD participant had two regions excluded (frontal and occipital).

### 2.3. Spectral power analysis

Spectral power was calculated separately for each channel and averaged across paired channels within each region of interest: F3/F4 (frontal), C3/C4 (central), and O1/O2 (occipital). Power spectra were computed using Welch’s method, applied to 2-second epochs with 50% overlap, resulting in a frequency resolution of 0.5 Hz. The final power spectra represent the median across all epochs. Absolute power values were normalized to relative power by dividing the power at each frequency by the total power across the analyzed range (1-50 Hz). Band-specific values were then computed by summing relative power values within the delta (2–4 Hz) and alpha (6–12 Hz) frequency bands. Absolute power values are presented in Figure S2; however, all analyses in this study were conducted using relative power. Relative power was analyzed, following standard conventions in developmental EEG research, particularly given the use of both clinical and high-density research systems with differing impedance properties (**Supplementary Table 3**).

### 2.3.1. Peak alpha frequency

Peak alpha frequency (PAF) was quantified separately for each of the six channels of interest using a custom MATLAB script designed to isolate periodic alpha activity (**Fig. 1a**) (Dickinson et al., 2018). To remove bias toward lower spectral frequencies, the aperiodic 1/f signal was first subtracted (Neto et al., 2015). To identify the peak within the alpha range (6–12 Hz), a Gaussian curve was fitted to the spectra using the MATLAB *fit* function (**Fig. 1b**). If the curve-fitting procedure failed, this was taken as evidence of no significant modulation in the alpha frequency range. These cases were independently reviewed by two authors (MBK, AHD) to confirm the absence of a peak. If power modulation was present but could not be accurately fitted due to irregular spectral curves, the peak was identified manually. After quantifying peaks at the channel level, we calculated regional values for each of the three regions by averaging the peak values across the two channels within the region. If a peak was detected in only one channel, that value was used as the regional value.

**Figure 1:**
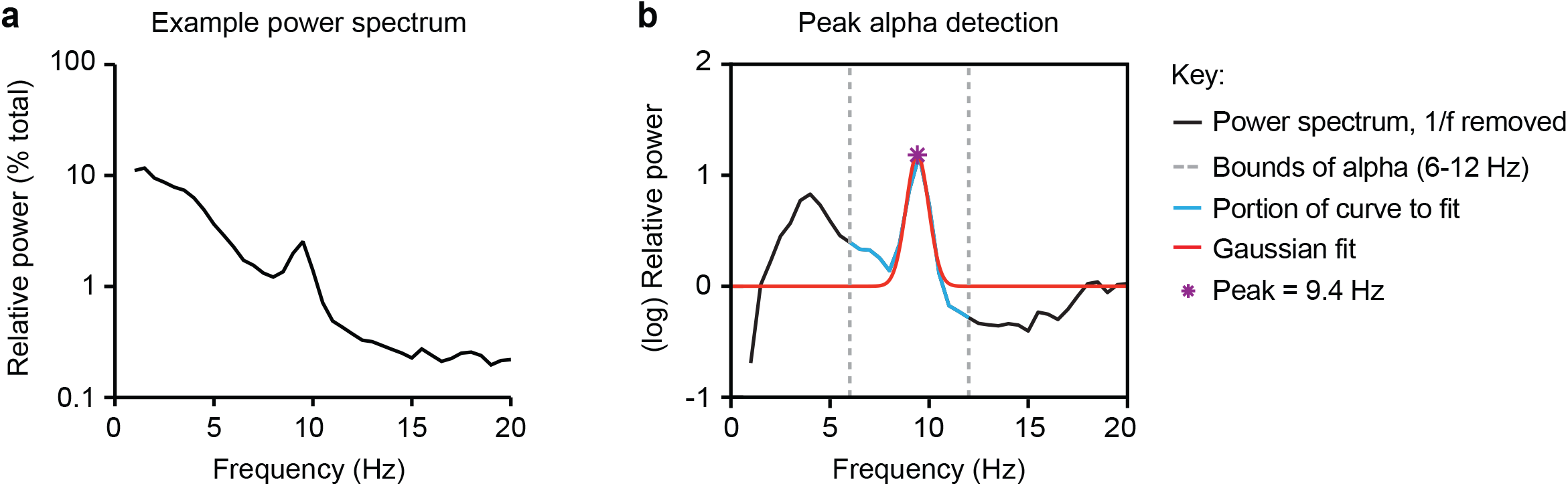
Methods for detecting peak alpha frequency (PAF). (a) Representative power spectrum from one channel in typically developing EEG, prior to removal of 1/f trend. (b) A Gaussian (red line) is fit to the 1/f-removed spectrum (black line) in the 6-12 Hz alpha range (blue line; range indicated by dotted gray line). The maximum of the Gaussian curve identifies the PAF.

### 2.3.2. Prominence threshold analysis

Given the low rates of peak detection in AS spectra, we explored an alternative peak detection method to determine whether these low rates were influenced by the detection approach itself. While Gaussian fitting is designed to identify distinct and well-defined spectral peaks, it may fail to capture subtler features. To address this limitation, we implemented MATLAB’s *findpeaks* function, which identifies peaks based on prominence thresholds. Prominence is defined as the vertical distance between the peak height and the average power of the nearest troughs on either side (see Fig. 7a). Peaks with greater prominence are more distinct from surrounding power. Prominence-based analysis was conducted on power values normalized to a 0–1 range to ensure consistency across participants, with 25 logarithmically spaced thresholds values ranging from 0.02 to 1. By adjusting the prominence threshold across this range of values, this method offered greater flexibility in identifying a wider range of spectral features, including subtle peaks that Gaussian fitting might miss.

### 2.4. Statistical analysis

Mixed-effects models were implemented to account for the hierarchical structure of the data, addressing non-independence due to some participants contributing multiple EEGs, and accounting for missing data as needed. Participant ID was included as a random effect in all models to account for repeated measures within participants.

### 2.4.1. Relative spectral power

Mixed-effects models were used to examine group differences in relative spectral power, specifically delta (2–4 Hz) and alpha (6–12 Hz) power, across frontal, central, and occipital regions. Fixed effects included group (AS vs. NT), age, and a Group × Age interaction, with participant ID included as a random intercept.

### 2.4.2. PAF

To investigate differences in peak detection rates, a generalized linear mixed-effects model (GLMM) with a logit link function was used, with fixed effects for group, region, and age, as well as their interactions. For participants with detectable peaks, a linear mixed-effects model was fitted to analyze differences in PAF. PAF values were modeled using mixed-effects models with group (AS vs. NT), age, and a Group × Age interaction as fixed effects, and participant ID included as a random intercept. Results are presented as mean ± SEM. Significance thresholds were defined as p<0.05, p<0.01, p<0.001, and p<0.0001, with corresponding annotations (*, **, ***, ****).

### 2.4.3. Prominence-based peak detection

We used a GLMM to examine the relationship between group (AS vs. NT), threshold, and their interaction on the likelihood of peak detection across varying thresholds (n=25). The model included fixed effects for group, threshold, and their interaction. At the median threshold value (0.14), we compared the primary Gaussian-based peak detection method with the secondary prominence-based analysis. Agreement in peak detection rates at this threshold was assessed using Cohen’s kappa. For participants with a detected peak at the median threshold, intraclass correlation coefficients (ICCs) were calculated to evaluate the consistency of peak frequency values between the two methods.

## 3. RESULTS

### 3.1. Increased delta power and decreased alpha power in AS

Linear mixed-effects models revealed significantly higher delta power (2–4 Hz) in Angelman syndrome (AS) compared to typically developing (TD) participants across all regions (Frontal: coefficient = -0.22, 95% CI [-0.26, -0.18], p < 0.0001; Central: coefficient = -0.246, 95% CI [-0.283, -0.209], p < 0.0001; Occipital: coefficient = -0.288, 95% CI [-0.324, -0.251], p < 0.0001; **Fig. 2a-b**). Relative delta power averaged across all electrodes is illustrated in **Fig. 2c**. Significant main effects of age indicated age-related declines in delta power for all regions (Frontal: coefficient = 0.00072, 95% CI [-0.0003, -0.001], p < 0.01; Central: coefficient = 0.00095, 95% CI [0.00052, 0.00139], p < 0.0001; Occipital: coefficient = 0.0011, 95% CI [0.00065, 0.00152], p < 0.0001). Additionally, significant Group × Age interactions (Frontal: coefficient = -0.001, 95% CI [-0.001, - 0.0007], p < 0.0001; Central: coefficient = -0.0013, 95% CI [-0.0016, -0.0010], p < 0.0001; Occipital: coefficient = -0.0017, 95% CI [-0.0021, -0.0014], p < 0.0001) showed that these age-related declines were more pronounced in AS compared to TD participants, suggesting steeper reductions in delta power with age in AS (results averaged across all electrodes illustrated in **Fig. 2d**).

**Figure 2:**
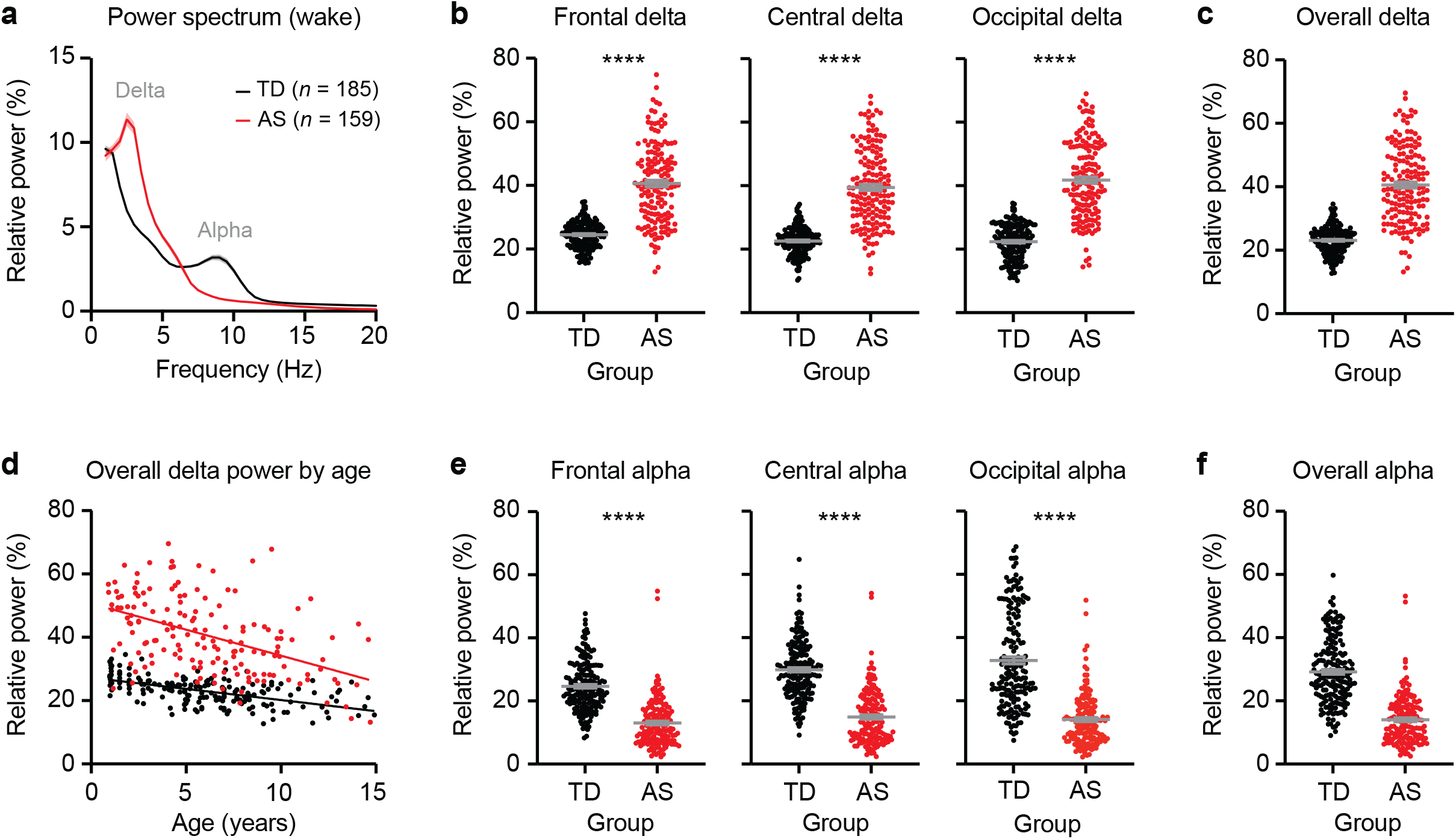
Children with AS have increased delta power and decreased alpha power. (a) Power spectrum of EEGs from typically developing (TD) children and children with AS during wake. Shaded area indicates ± SEM. (b) Relative delta power (2-4 Hz) is increased in AS in frontal, central, and occipital channels. Frontal: TD: *n* = 184, AS: *n* = 156; Central: TD: *n* = 185, AS: *n* = 159; Occipital: TD: *n* = 183, AS: *n* = 157. (c) Relative delta power, averaged across all electrodes. (d) Relative delta power as a function of age in TD and AS EEGs. (e) Relative alpha power (6-12 Hz) is decreased in AS in frontal, central, and occipital channels. (f) Relative alpha power, averaged across all electrodes. Data plotted as mean ± SEM; \*\*\*\**p* < 0.0001.

Linear mixed-effects models indicated significantly lower relative alpha power (6–12 Hz) in AS compared to TD participants across all three regions (Frontal: coefficient = 0.133, 95% CI [0.103, 0.162], p < 0.0001; Central: coefficient = 0.184, 95% CI [0.149, 0.218], p < 0.0001; Occipital: coefficient = 0.159, 95% CI [0.111, 0.207], p < 0.0001; **Fig. 2e**). Relative alpha power averaged across all electrodes is illustrated in **Fig. 2f**. Significant main effects of age revealed that alpha power increased with age in both groups (Frontal: coefficient = 0.00083, 95% CI [0.00059, 0.00107], p < 0.0001; Central: coefficient = 0.00104, 95% CI [0.00078, 0.00130], p < 0.0001; Occipital: coefficient = 0.00117, 95% CI [0.00088, 0.00145], p < 0.0001). Group x Age interactions indicated that age-related changes in alpha power were comparable across the two groups in frontal and central regions, with a significant Group X Age interaction occipitally (Frontal: coefficient = -0.0001, 95% CI [-0.00044, 0.00024], p = 0.565; Central: coefficient = -0.00029, 95% CI [-0.00068, 0.00010], p = 0.143; Occipital: coefficient = 0.00062, 95% CI [0.00010, 0.00114], p = 0.020). Spectral analyses of raw delta and alpha power are reported in **Fig. S2**.

### 3.2. Reduced PAF presence in AS

In the AS group, 39.74% of participants showed an alpha peak in the frontal region, 45.28% in the central region, and 42.41% in the occipital region. Comparatively, in the TD group, 92.39% showed a peak in the frontal region, 95.14% in the central region, and 94.57% in the occipital region (**Fig. 3a**). When considering participants who showed a peak in any region, 59.75% of the AS group displayed a peak, compared to 95.68% in the TD group (**Fig. 3b**). To formally compare rates of alpha peak presence, we focused on regional-level data. A logistic mixed-effects model revealed a significant main effect of group (p < 0.0001), with TD participants being more likely to exhibit a peak compared to AS participants (log-odds = 3.75, 95% CI [2.48, 5.02]). A significant main effect of age (p < 0.0001) indicated that peak presence increased with age (log-odds = 0.019, 95% CI [0.012, 0.027]). The non-significant Group × Age interaction (p = 0.76) suggests that this age-related increase was consistent across both groups. No significant differences in peak presence were observed between regions (p > 0.05), and Group × Region interactions were also non-significant (p > 0.05).

**Figure 3:**
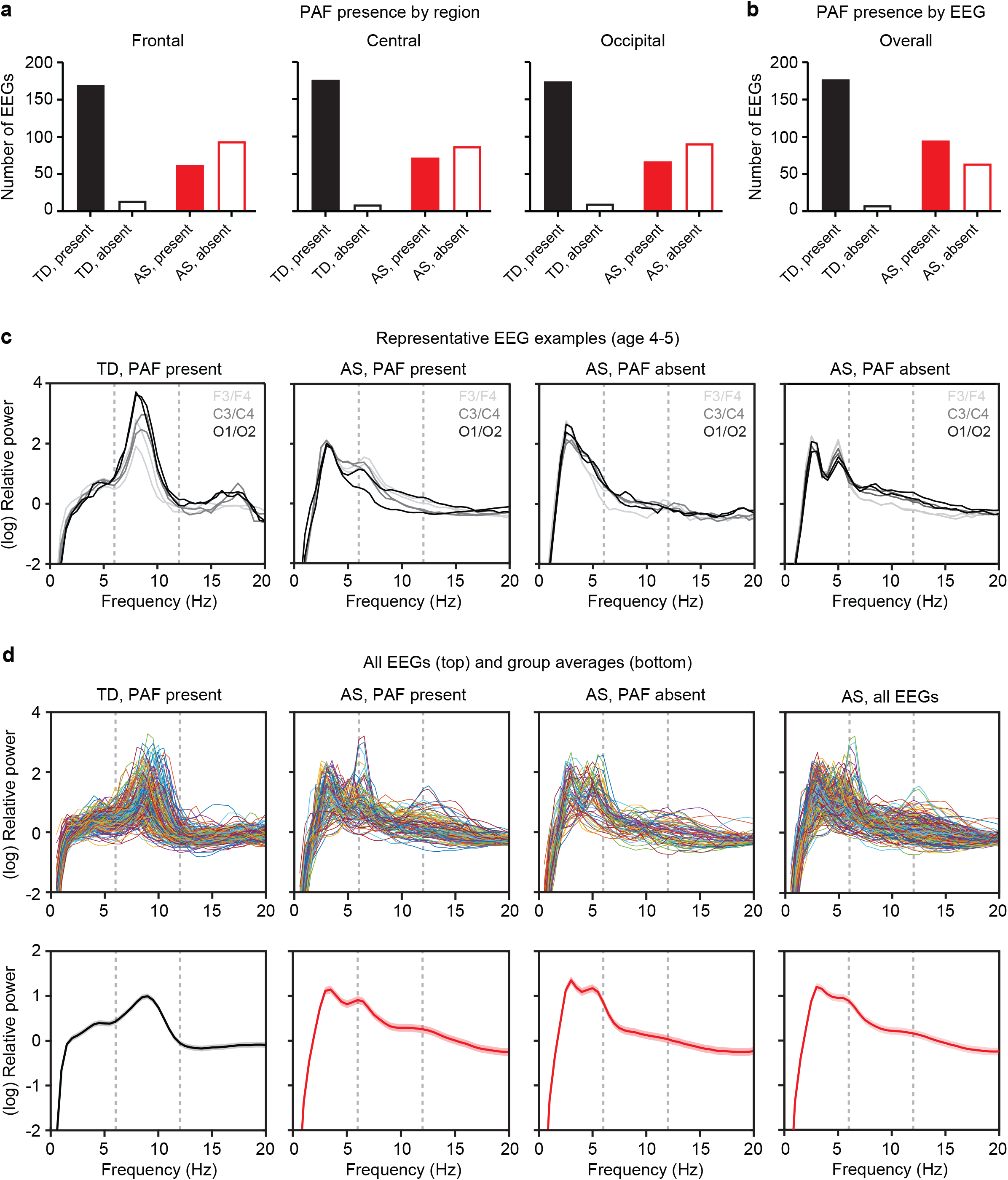
PAF was not present in a subset of AS EEGs. (a) PAF presence by region in TD and AS EEGs. (b) PAF presence by EEG in TD and AS EEGs. (c) Representative 1/f-removed spectra from four subjects illustrate typical detection of PAF in a TD EEG (left), qualitatively abnormal PAF in an AS EEG (second from left), absence of PAF in an AS EEG without (second from right) and with a theta peak outside the range of alpha (right). Six traces represent six channels in each plot. (d) Top panels show 1/f-removed spectra from all EEGs and bottom panels show group averages; shaded area indicates ± SEM. Dotted lines represent bounds of alpha (6-12 Hz).

Figure 3 also illustrates 1/f-removed spectra across channels for four representative EEGs (1 TD; 3 AS; **Fig. 3c**), and individual traces for each EEG (**Fig. 3d**).

### 3.3. Atypical developmental PAF trajectory in AS

In EEGs where an alpha peak was present, linear mixed-effects models revealed a significant main effect of group in central and occipital but not frontal regions (Frontal: coefficient = 0.463, 95% CI [-0.184, 1.111], p = 0.159; Central: coefficient = 0.708, 95% CI [0.202, 1.213], p < 0.01; Occipital: coefficient = 0.755, 95% CI [0.252, 1.258], p < 0.01), with TD participants showing higher PAF values compared to AS participants (**Fig. 4a**). PAF averaged across all electrodes is illustrated in **Fig. 4b-c**. There was a significant main effect of age indicating an age-related increase in PAF for the occipital but not frontal or central region (Frontal: coefficient = 0.0151, 95% CI [-0.056, 0.0863], p = 0.675; Central: coefficient = 0.00638, 95% CI [-0.0494, 0.0622], p = 0.8220; Occipital: coefficient = 0.0761, 95% CI [0.0208, 0.1314], p < 0.01). Significant Group × Age interactions in all regions indicated that TD participants exhibited significantly steeper age-related PAF increases (Frontal: coefficient = 0.1508, 95% CI [0.0689, 0.232], p < 0.001; Central: coefficient = 0.1787, 95% CI [0.1134, 0.2439], p < 0.0001; Occipital: coefficient = 0.1501, 95% CI [0.0844, 0.2157], p < 0.0001; **Fig. 4d**). **Fig. 5** illustrates individual 1/f-removed spectra for all EEGs, categorized by age and genotype.

**Figure 4:**
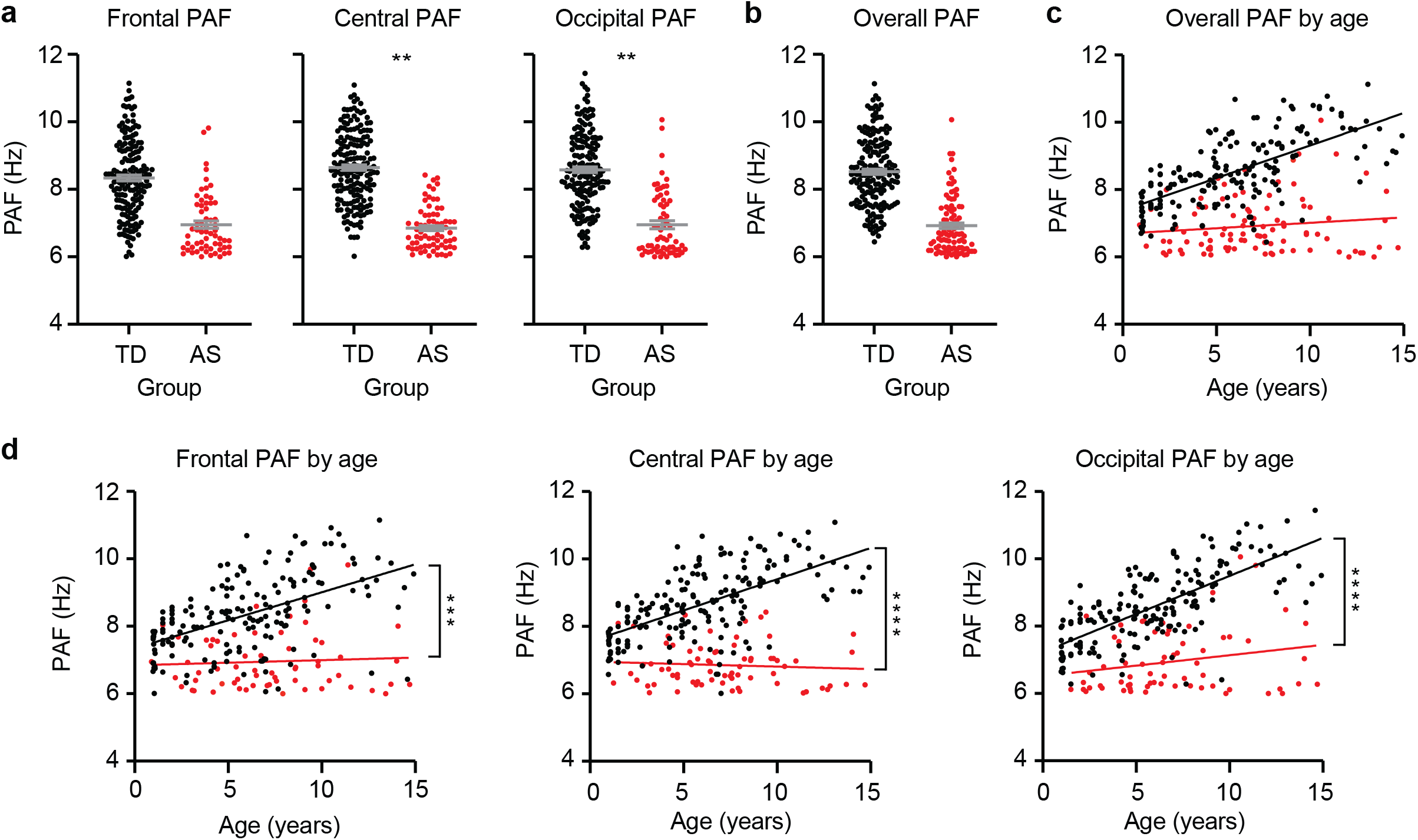
PAF does not develop normally in AS EEGs. (a) Comparison of PAF by genotype in frontal, central, and occipital regions. (b) Average PAF across all channels and (c) as a function of age. (d) PAF as a function of age in frontal channels, central channels, and occipital channels. Each point represents the averaged PAF of two channels per region of interest per subject. Error bars indicate ± SEM; \*\**p* < 0.01, \*\*\**p* < 0.001, *\*\*\*\*p* < 0.0001. Asterisks in (d) represent Genotype X Age interaction.

**Figure 5:**
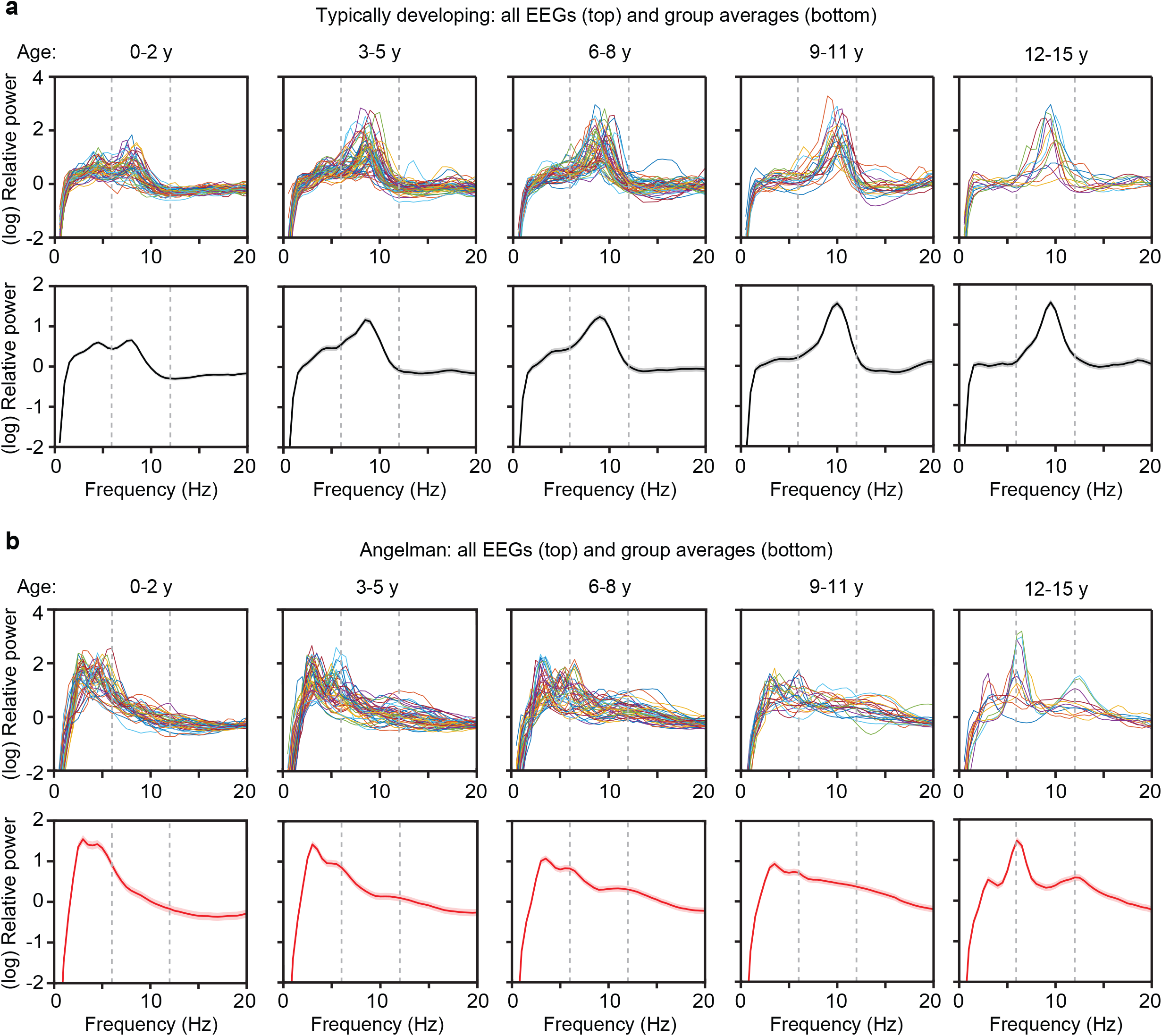
PAF development by age. (a) Top panels illustrate 1/f-removed spectra from all TD EEGs (averaged across channels) for each age group; 0-2 years old (*n* = 48, left), 3-5 years old (*n* = 54, second from left), 6-8 years old (*n* = 48, middle), 9-11 years old (*n* = 23, second from right), 12-15 years old (*n* = 12, right). Bottom panels illustrate averages for each age group; shaded area indicates ± SEM. (b) Top and bottom panels illustrate 1/f-removed spectra from all AS EEGs and group averages by age (0-2: *n* = 37, 3-5: *n* = 48, 6-8: *n* = 39, 9-11: *n* = 23, 12-15: *n* = 12).

### 3.4. Validation of reduced peaks and PAF using prominence analyses

Given the low rates of peak detection in AS spectra (**Fig. 3a-b**), we explored an alternative prominence-based peak detection method (**Fig. 6a**) to determine whether these low rates were influenced by Gaussian fitting process. Peak detection rates were consistently lower in AS participants compared to TD participants across all thresholds, including the most lenient (**Fig. 6b**). A linear mixed effects model demonstrated a significant effect of group (-3.8443, p < 0.001), indicating that participants in the AS group have significantly lower odds of peak detection compared to TD participants, and a significant effect of threshold (-12.389, p < 0.001), indicating that increasing threshold values are strongly associated with reduced odds of detecting a peak across all participants. The interaction effect between group and threshold (-0.0204, p = 0.973) was not statistically significant, indicating that the effect of threshold on peak detection does not differ significantly between AS and TD participants. This suggests that the absence of peaks in AS spectra was not due to the Gaussian fitting process being overly strict or failing to detect less distinct peaks in AS.

**Figure 6:**
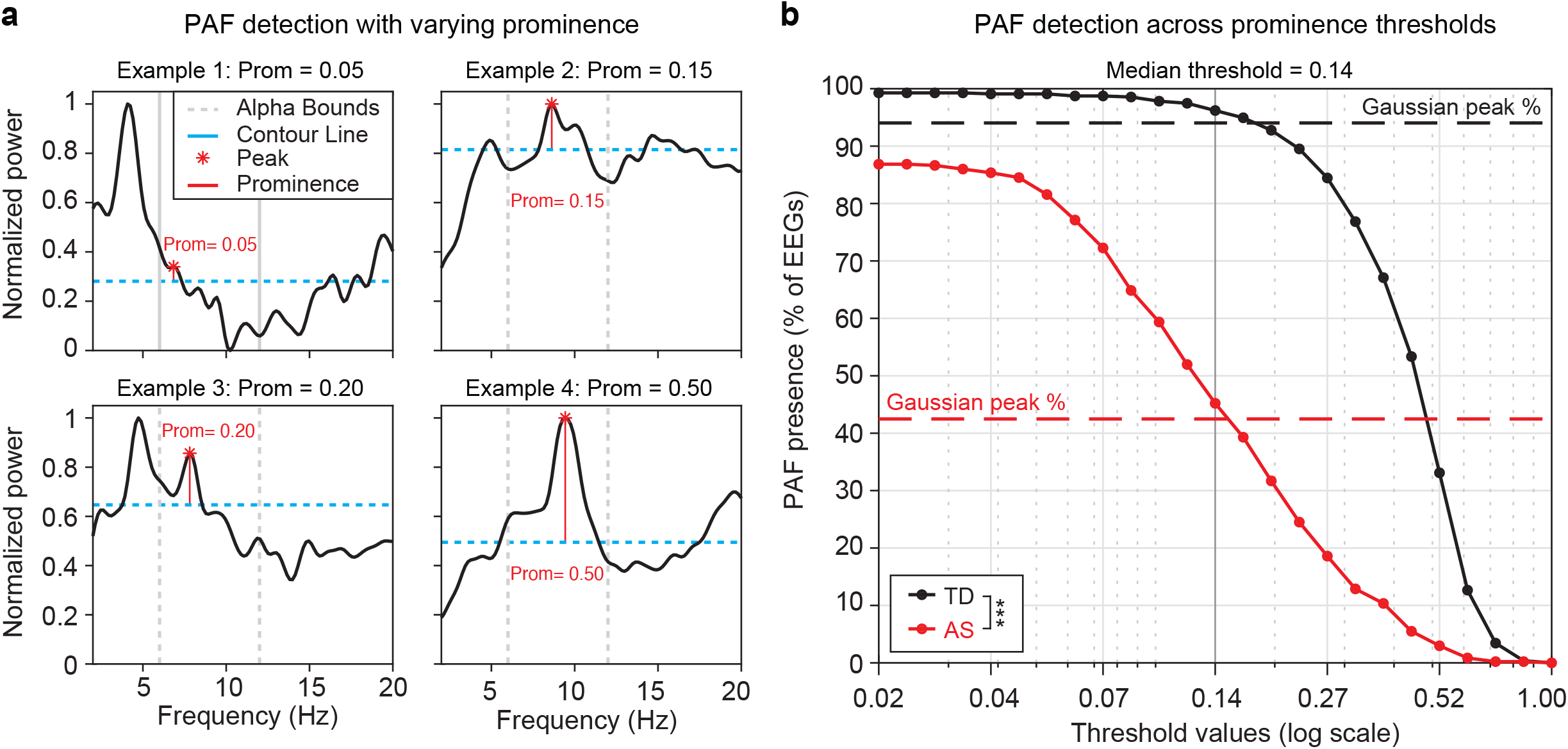
Prominence detection confirms abnormal alpha oscillations in Angelman EEGs. (a) Example spectra illustrating peaks at varying prominence levels. (b) Percentage of peaks detected in AS participants (red) vs. TD participants (black) across different threshold values. Dashed vertical lines represent the percentage of peaks detected for each group using Gaussian analysis, while the solid black vertical line indicates the median threshold (0.14).

Peak detection rates at the median threshold (0.14) were 69.41%, 76.45%, and 72.22% for the frontal, central, and occipital regions, respectively. These rates were consistent with those obtained using the Gaussian approach (68.24%, 72.09%, and 70.47%). Cohen’s kappa statistics demonstrated substantial inter-rater agreement in peak detection rates between the two methods: frontal (κ = 0.70, 95% CI [0.603, 0.767]), central (κ = 0.611, 95% CI [0.516, 0.696]), and occipital (κ = 0.61, 95% CI [0.522, 0.70]). For participants with a detected peak, PAF values were highly consistent between the two methods, with average differences of -0.38 Hz (frontal), -0.31 Hz (central), and -0.34 Hz (occipital). Intraclass correlation coefficients (ICCs) further confirmed the strong agreement between methods, with high consistency observed across all regions: ICC = 0.86 (frontal), ICC = 0.90 (central), and ICC = 0.78 (occipital), all p < 0.0001.

## 4. DISCUSSION

Alpha oscillations undergo well-characterized developmental changes that closely align with neural circuit maturation, making them a promising focus for understanding atypical brain function. This study provides novel insights into alpha metrics, including PAF, to determine their value as markers for characterizing and mapping atypical neuronal dynamics in AS. Our results demonstrate that PAF is highly sensitive to loss of neuronal Ube3a expression, providing a potential marker to better understand both the underlying neuropathology and the unique developmental trajectories associated with AS.

Consistent with previous findings, we observed higher delta power and lower relative alpha power in children with AS (**Fig. 2**), indicating altered oscillatory dynamics. To further probe these dynamics, we quantified PAF and found that TD children consistently displayed pronounced alpha peaks and exhibited typical age-related increases in PAF. In contrast, developmental patterns were significantly disrupted in AS. Children with AS were significantly less likely to exhibit an alpha peak compared to age-matched typically developing controls (**Fig. 3**), and among those who did show an alpha peak, PAF was significantly lower (**Fig. 4**). Furthermore, while prominent in the TD group, the well-established age-related increase in PAF was absent in AS (**Figs. 4-5**).

In addition, when PAF was observed in AS EEGs, the shape of 1/f-removed spectra appeared qualitatively different than in TD controls (**Fig. 3c-d**). The TD subjects consistently displayed clear, pronounced peaks in the alpha band, whereas PAF was decreased in both amplitude and frequency and appeared qualitatively “immature”, resembling spectra often seen in typically developing infants (**Fig. 3c**) (Marshall et al., 2002; Wilkinson et al., 2024). The examples shown in **Fig. 3c** are representative, and the less mature alpha profile consistent with earlier developmental stages was also observed across group averages of AS EEGs where PAF was determined to be present (**Fig. 3d**). Even when using a more sensitive prominence-based peak detection method, AS participants consistently exhibited lower peak rates compared to TD children, regardless of the threshold applied (**Fig. 6**). These consistent findings across detection methods suggest that the atypical alpha dynamics in AS reflect genuine neurophysiological differences rather than methodological artifacts.

### 4.1. Implications for alpha metrics in AS

These results highlight the potential of alpha oscillations, specifically the presence and frequency of an alpha peak, as critical markers for assessing brain function in AS. Specifically, alpha metrics are highly sensitive to atypical circuit function, making them potentially valuable for tracking responses to treatments aimed at improving brain function (Dickinson et al., 2018; Edgar et al., 2019). When combined with other measures, such as delta power, alpha metrics may provide a powerful framework for detecting changes in neural circuits targeted by disease-modifying treatments, offering insights that precede observable downstream behavioral improvements. For instance, alpha metrics may directly index maturational changes in the underlying large-scale circuits that support increasingly complex cognition and behavior (Klimesch et al., 2007; Marshall et al., 2002). Such early detection capabilities are critical for evaluating intervention efficacy and may help address the critical need for quantifiable biomarkers to support and enhance clinical endpoints.

Furthermore, these EEG-based measures offer a unique opportunity to serve as translational biomarkers, providing a consistent framework for evaluating neural changes across both preclinical and clinical trials. This dual utility underscores their promise as tools for advancing therapeutic development and understanding the mechanisms underlying treatment effects in AS.

Alpha metrics may also complement delta metrics by providing distinct and complementary insights into neural function. Given that elevated delta power reflects generalized circuit disruption, therapeutic improvements in delta power (i.e., reductions) would likely reflect circuits becoming less disorganized and movement towards more typical circuit dynamics. This aligns with the observed age-related reductions in delta power, which may represent the natural progression of neural circuits maturing and transitioning toward more efficient states. In contrast, alpha metrics, such as PAF, may offer a finer-grained measure of how circuits are actively maturing and achieving greater functional specialization. Furthermore, because alpha metrics undergo rapid and measurable changes during early development as circuits mature (Dickinson et al., 2025; Wilkinson et al., 2024), they may be particularly well-suited for detecting fast, subtle changes in neural activity over the course of a clinical trial. PAF, in particular, is a valuable metric because it remains generally stable across behavioral conditions and is quantified in a way that is not influenced by increased delta (Dickinson et al., 2018; Grandy et al., 2013; Haegens et al., 2014; Petrosino et al., 2018). This sensitivity to developmental dynamics positions alpha metrics as powerful tools for tracking treatment effects and understanding neural mechanisms underlying variable trajectories in AS.

### 4.2. Limitations and next steps

While these findings underscore the significant potential of alpha metrics as sensitive markers of neural function and maturation, this study is not without limitations, and important next steps are needed to advance our understanding of alpha metrics in this context. One potential limitation is that EEGs from children with AS were collected using clinical systems, whereas EEGs from neurotypical controls were obtained from high-density research systems. Although this difference could introduce small variations in signal characteristics, the stark differences observed between the AS and NT groups are unlikely to be solely attributable to equipment variations. Additionally, both clinical and research systems are widely validated for capturing reliable neural signals in pediatric populations. Future studies with identical EEG systems across groups would further ensure methodological consistency; however, the pronounced findings here strongly suggest that the observed atypical alpha dynamics in AS reflect a true underlying neurophysiological difference rather than an artifact of system differences.

Another limitation of this study is that the age-related changes described are based on cross-sectional data, which provide a snapshot of developmental trends across different individuals but may not fully capture the dynamic trajectories of neural changes over time. Future work may leverage data from individuals with multiple data points to provide a more comprehensive report of longitudinal developmental PAF patterns in AS and how these changes map onto variations in cognitive function, and changes over time. In addition, while this study highlights alpha metrics as markers of neural maturation, further work is needed to investigate how these neural changes map onto cognitive development. Understanding the relationship between alpha oscillations and cognitive function is crucial for determining whether alpha metrics can serve as meaningful biomarkers of functional outcomes. This would provide critical context for interpreting their potential role in tracking developmental changes and assessing the efficacy of interventions in AS.

Finally, as an initial characterization of alpha metrics, this study used the 6–12 Hz range to examine alpha dynamics, consistent with frequency ranges commonly applied in developmental populations. However, given the dysmaturation of alpha oscillations observed in our analyses, it is important to recognize the limitations of using fixed frequency boundaries during early spectral maturation. In typical development, spectral maturation involves a gradual reorganization of power across frequencies (Dickinson et al., 2025; Patino et al., 2017; Xiao et al., 2015). Therefore, adopting a developmental framework that captures these broader shifts in power may offer a more flexible and accurate approach for quantifying spectral dynamics in AS, which deviates significantly from age-expected patterns of change. Future work should explore frequency ranges that are not strictly predefined, enabling a more comprehensive and sensitive assessment of neural maturation and atypical development in AS. A broad-spectrum marker would continuously capture evolving oscillatory dynamics, providing a robust tool for monitoring developmental and treatment effects across varied stages of maturation.

## Acknowledgments

We thank Katherine Walsh (MGH) and Anne Wheeler (RTI) for their support in collating data.

## Author Contributions

Conceptualization: AHD & MSS, data curation: AHD, MBK, JJS, AEY, CAM, DPR, CJC, MSS, formal analysis: AHD, MBK, JJS, AEY, MSS, funding acquisition: AHD & MSS, methodology: AHD, MBK, MSS, software: AHD, NSH, MSS, supervision: AHD & MSS, validation: AHD, MBK, MSS, visualization: AHD, MBK, JJS, MSS, writing – original draft: AHD, MBK, MSS, writing – review and editing: all authors.

## FIGURE LEGENDS

**Figure S1:**
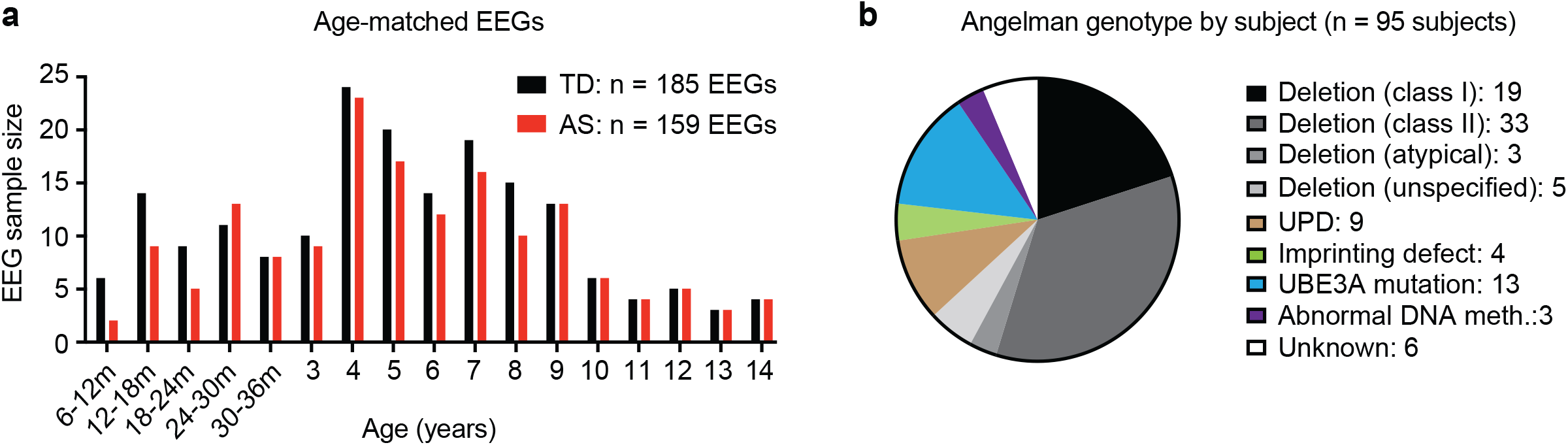
Age and genotype information. (a) 159 AS EEGs from 96 subjects were age-matched to 185 EEGs from typically developing (TD) subjects. (b) AS genotype.

**Figure S2:**
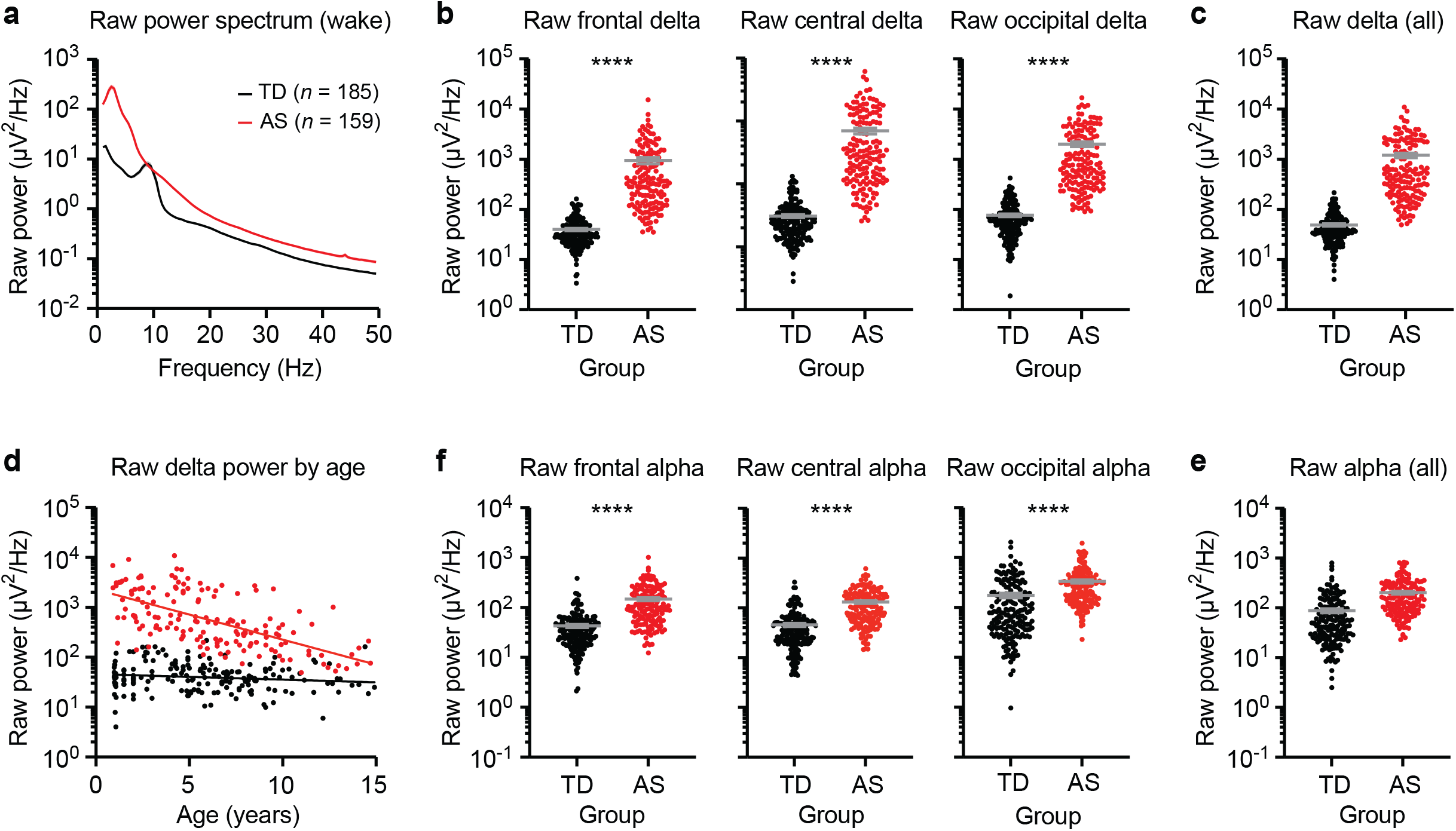
Raw spectral power is increased in AS EEGs across bands. (a) Raw power spectrum of EEGs from typically developing (TD) children and children with AS during wake. Shaded area indicates ± SEM. (b) Raw delta power (2-4 Hz) is increased in AS EEGs in all regions (Frontal: coefficient = -1748.9, 95% CI [-2240.4, -1257.4], p < 0.0001; Central: coefficient = -1418.3, 95% CI [-1675.2, -1161.4], p < 0.0001; Occipital: coefficient = -4231.2, 95% CI [-4959.4, -3503.1], p < 0.0001). Frontal: TD: *n* = 184, AS: *n* = 156; Central: TD: *n* = 185, AS: *n* = 159; Occipital: TD: *n* = 183, AS: *n* = 157. (c) Raw delta power averaged across all electrodes. (d) Raw delta power (averaged across all electrodes) as a function of age in TD and AS. (e) Raw alpha power (6-12 Hz), is increased in AS EEGs in all regions (Frontal: coefficient = -159.6, 95% CI [-201.3, -117.9], p < 0.0001; Central: coefficient = -149.5, 95% CI [-181.9, -117.0], p < 0.0001; Occipital: coefficient = -396.3, 95% CI [-517.1, -275.5], p < 0.0001). (f) Raw alpha power averaged across all electrodes. Data plotted as mean ± SEM; \*\*\*\**p* < 0.0001.

