## Supplement for "Atypical Alpha Oscillatory EEG Dynamics in Children with Angelman Syndrome"

| Number EEGs Contributed | Number of participants |
| --- | --- |
| 1 EEG | 60 |
| 2 EEGs | 16 |
| 3 EEGs | 12 |
| 4 EEGs | 5 |
| 5 EEGs | 1 |
| 6 EEGs | 1 |

**Supplementary Table 1.** Number of AS EEGs contributed per participant.

|  | **AS EEGs** | |
| --- | --- | --- |
|  | *Site 1 (UNC)* | *Site 2 (ASNHS)* |
| N EEGs | 10 | 149 |
| Age (y) | 5.1 ± 1.2 | 6.2 ± 0.3 |
| Sex | M: 5 (50%)  F: 5 (50%) | M: 97 (65%)  F: 52 (35%) |
| EEG system | Natus or Grass | Bio-Logic or Xltek |
| N Channels | 6 (F3, F4, C3, C4, O1, O2) | 10-20 system |
| Sampling rate (Hz) | 200 or 256 | 200, 256, or 512 |
| Recording length (min) | 76.8 ± 18.5 | 21.2 ± 1.5 |
| Reference | Vertex | Varied |

**Supplementary Table 2.** Site-by-site properties of AS EEGs.

|  | **TD EEGs** | | |
| --- | --- | --- | --- |
|  | *Site 1 (UNC)* | *Site 2 (MGH)* | *Site 3 (UCLA)* |
| N EEGs | 10 | 98 | 77 |
| Age (y) | 5.8 ± 1.3 | 7.5 ± 0.3 | 3.7 ± 0.3 |
| Sex | M: 5 (50%)  F: 5 (50%) | M: 52 (53%)  F: 46 (47%) | M: 42 (55%)  F: 35 (45%) |
| EEG system | Natus or Grass | Natus | EGI |
| N Channels | 6 (F3, F4, C3, C4, O1, O2) | 10-20 system | 129-channel HydroCel Geodesic Sensor net |
| Sampling rate (Hz) | 200 or 256 | 200, 250, 256, 500, or 512 | 250 |
| Recording length | 67.3 ± 12.3 | 10.6 ± 1.3 | 2.5 ± 0.2 |
| Reference | Vertex | C2 spinous process | Vertex |

**Supplementary Table 3.** Site-by-site properties of TD EEGs.
